# Alternative translation initiation generates a functionally distinct isoform of the stress-activated kinase MK2

**DOI:** 10.1101/429696

**Authors:** Philipp Trulley, Goda Snieckute, Dorte Bekker-Jensen, Manoj B. Menon, Robert Freund, Alexey Kotlyarov, Jesper V. Olsen, Manuel D. Diaz-Muñoz, Martin Turner, Simon Bekker-Jensen, Matthias Gaestel, Christopher Tiedje

**Affiliations:** Institute of Cell Biochemistry, Hannover Medical School (MHH), Carl-Neuberg-Str. 1, D-30625 Hannover, Germany; Center for Healthy Aging, Department of Cellular and Molecular Medicine, University of Copenhagen, Blegdamsvej 3B, DK-2200 Copenhagen N, Denmark; Mass Spectrometry for Quantitative Proteomics, Proteomics Program, The Novo Nordisk Foundation Center for Protein Research, Faculty of Health and Medical Sciences, University of Copenhagen, Blegdamsvej 3B, DK-2200 Copenhagen N, Denmark; Centre de Physiopathologie Toulouse-Purpan, INSERM UMR1043/CNRS U5282, Toulouse, 31300, France; Lymphocyte Signalling and Development, The Babraham Institute, CB22 3AT, Cambridge, United Kingdom

**Keywords:** ribosome footprinting, alternative translation initiation, 5′UTR, eIF4A1, MAPKAPK2, mass-spectrometry

## Abstract

Shaping of the proteome by alternative translation is an important mechanism of post-transcriptional gene regulation. It can lead to the expression of multiple protein isoforms originating from the same mRNA. Here we show that a novel, abundant and long isoform of the stress/p38^MAPK^-activated kinase MK2, a key regulator of transcription, migration, death signaling and post-transcriptional gene regulation, is constitutively translated from an alternative CUG translation initiation start site located in the 5′UTR of its mRNA. GC-rich sequences and putative G-quadruplex structures influence the usage of that codon as a translation initiation start site and the RNA helicase eIF4A1 is needed to ensure alternative isoform translation. We recapitulated the usage of the alternative start codon and determined the molecular properties of the short and a long MK2 isoforms. Phenotypically, only the short isoform phosphorylated Hsp27, supported migration and stress-induced immediate early gene (IEG) expression. Interaction profiling by quantitative mass-spectrometry revealed short isoform-specific binding partners that were associated with migration. In contrast, the long isoform contains additional putative phosphorylation sites in its unique N-terminus. In sum, our data reveal a longer and previously non-described isoform of MK2 with distinct physiological properties originating from alternative translation.

## Introduction

Post-transcriptional gene regulation (PTGR) constitutes an important toolbox used to shape gene expression upon intra- and extracellular cues. PTGR impacts on mRNA processing, localization, export, stability and translation. Canonical translation of mRNAs starts on AUG translation initiation sites (TIS) that are recognized by ribosomes through scanning of the 5’ UTR leader sequence and is dependent on the cap-structure. Translation initiation and the production of protein isoforms by alternative translation mechanisms are highly regulated. They are influenced by sequences in the 5’ untranslated region (5′UTR) bearing internal ribosomal entry sites (IRES), secondary structures, upstream open reading frames (uORFs) or by “leaky scanning” of sequences preceding the initiator AUG (Kozak sequence), leading to the usage of alternative start codons (1,2). Additionally, numerous translation events at non-AUG start codons have been detected in mRNAs determining alternative translation initiation sites (aTIS) sequences (3,4). The mechanisms by which ribosomes select non-AUG codons for translation initiation might rely on uORFs, structured 5′UTRs (5) or translation initiation auxiliary factors and RNA helicases such as eIF4A1, eIF2A or Ddx3 (6–8). Taken together these features of translation explain in part how proteome diversity in cells can be generated from a limited number of transcripts.

Protein kinases and associated signal transduction pathways play an important role in regulating transcriptional and post-transcriptional events. A well-characterized pathway is the p38^MAPK^-MK2-signaling module. The p38^MAPK^-activated kinase MK2 (or MAPKAPK2) phosphorylates substrates such as the RNA-binding protein and PTGR master regulator tristetraprolin (TTP) and the transcription factor serum response factor (SRF) (9). Numerous reports characterize the diverse functions of MK2 in inflammation and stress signaling (10), also suggesting its role as a checkpoint kinase in cancer (11). Powerful tools for these studies are the MK2 knockout (MK2 KO) (12) and the MK2/3 double knockout (MK2/3 DKO) (13) mouse models and, consequently, many phenotypes could be detected with the help of these animals. These phenotypes range from defects in immediate early gene responses over impaired cytokine production and regulation of apoptosis/necroptosis to non-coding RNA regulation and defects in migration. The above mentioned phenotypes could be linked to specific substrates of MK2, such as the above mentioned TTP (14) and SRF (15), RIPK1 (16–18) and RBM7 (19,20).

Western blot detection of endogenous MK2 often gave two bands where only the lower band represented the apparent molecular mass of the kinase (21,22). Interestingly, ectopically expressed MK2 does not display the second, slower migrating band. To date, no information about the nature of this MK2 isoform expression is available. By exploiting RNA sequencing (RNASeq), ribosome footprinting (RiboSeq) and by conducting biochemical assays, we show that the long MK2 isoform constitutively arises from alternative translation of an in-frame upstream CUG alternative translation initiation site (aTIS) in the highly structured and conserved GC-rich 5′UTR. The CUG is surrounded by an alternative Kozak sequence, and the helicase eIF4A1 is needed to secure translation of both the CUG- and AUG-initiated isoforms. Mechanistically, the long and short MK2 isoforms engage in distinct protein-protein interactions and rescue MK2-deficiency phenotypes to different extents *in vivo*. Therefore, our results resolve the controversy surrounding an additional high molecular weight MK2 band often noted since its discovery in the 1990s (21,22).

## Results

### Translation of MK2 from an alternative start codon in the 5′UTR of its mRNA

MK2 is a ubiquitously expressed p38^MAPK^-activated protein kinase (10). When analyzing its expression in different cell lines of murine and human origin or in mouse tissues by Western blotting, a multiple band pattern is detected (Fig. 1A,B, S1). Of note, the ratio of isoform expression is distinct and the intensities of the lower and upper band vary between tissues and cell lines. In murine cells and tissues, two predominant bands of approximately 50 and 60 kDa can be detected in Western Blot by using an anti-MK2 antibody directed towards the C-Terminus of human MK2 (Fig. 1A,B). The low molecular weight isoform corresponds to the hitherto known open reading frame (ORF) of the gene, whereas the origin and function of the slower migrating isoform is obscure, and only one transcript existed in the NCBI database (NM_008551.2). Importantly, both bands were found to be absent in cells from MK2 knockout (KO, described in (12)) and MK2/3 double knockout mice (DKO, described in (13)) (Fig. 1A, S1A). In human cell lines two dominating bands of around 55 kDa together with two less intense bands at around 65 kDa were detected (Fig. 1A, S1C,D). Two similar human MK2 transcripts (NM_004759.4 and NM_032960.3) can be found in the database and arise from alternative splicing. shRNA-mediated knockdown of human MK2 in HT29 cells removed all of these bands confirming that they were derived from the same gene (Fig. S1D). For the murine MK2-related protein kinase MK3 only one specific protein product is detected and this is absent in MK2/3 DKO cells (Fig. S1A,B). Taken together, we concluded that the different MK2 protein bands can be assigned to distinct protein isoforms that may originate from post-translational modifications of MK2, alternative splicing or alternative translation events.

**Figure 1.**
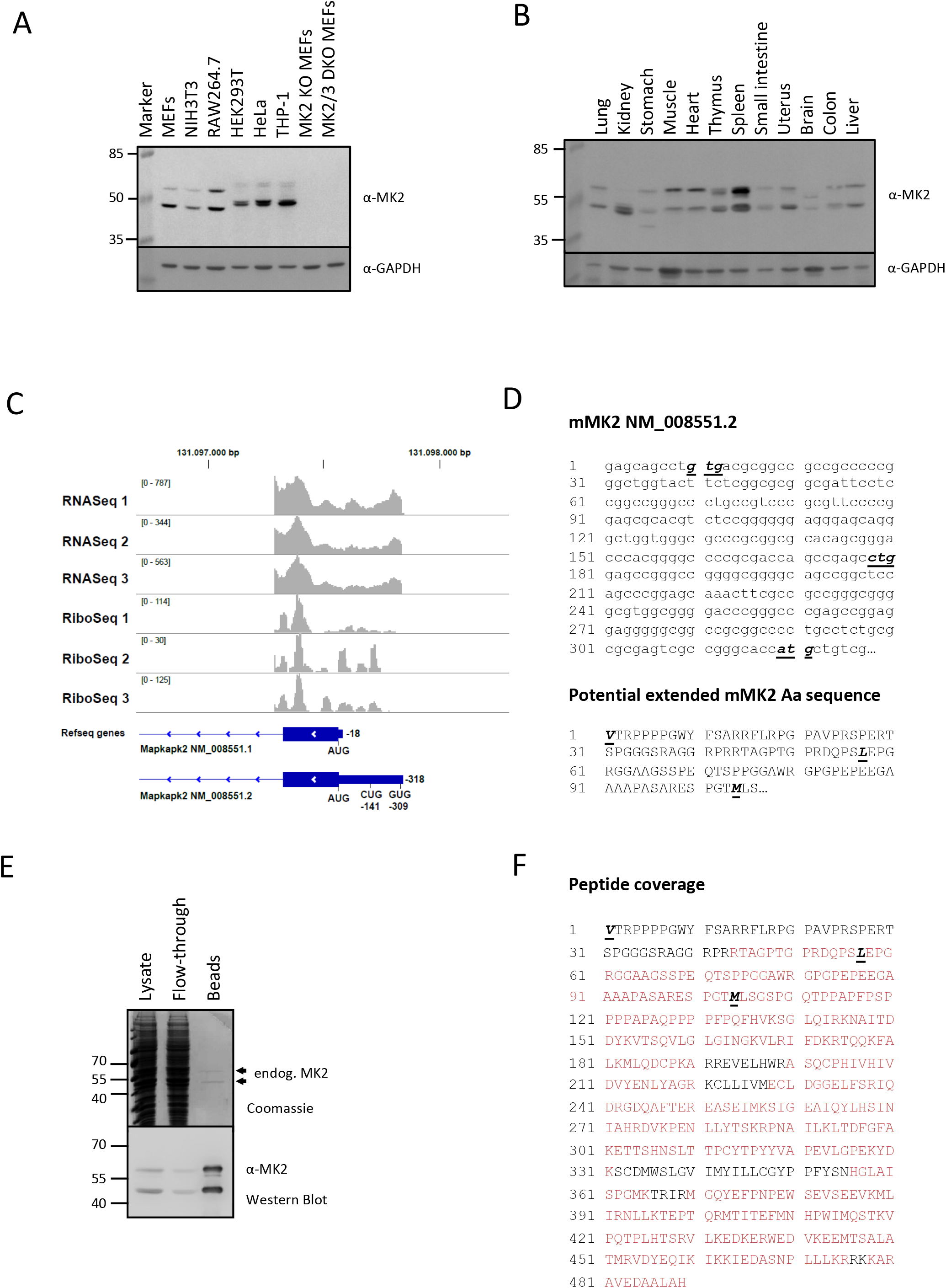
Translation of MK2 from an alternative start codon in its 5′UTR. **A**. Expression profiling of MK2 in different murine and human cell lines. Lysates from the indicated cell lines were subjected to Western Blot analysis using an anti-MK2 antibody (D1E11, Cell Signaling) and an anti-GAPDH as loading control. **B**. Expression of MK2 in several mouse organs as detected by Western Blot analyses using a MK2-specific and a MK3-specific antibody. The GAPDH development served as a loading control. **C**. Genomic tracks of RNASeq and RiboSeq data sets (GSE81250) for the old murine MK2 transcripts NM_008551.1 and its newest annotated transcript version NM_008551.2. The position of potential alternative translation initiation sites (aTIS) within the 5′UTR of MK2 and the AUG initiation sites are indicated. The length of both 5′UTR are given (18 and 318 nucleotides, respectively). **D**. The murine MK2 5′UTR nucleotide sequence and its putative extended in-frame N-terminal amino acid sequence. The aTIS are marked in bold and in italics within both nucleotide and the amino acid sequence. **E**. Purification of endogenous MK2 isoforms form RAW264.7 cells. Endogenous MK2 was purified from 1 × 10^8^ RAW264.7 cells using an agarose MK2-Trap (Chromotek). Input (lysate), column flow-through and precipitates (beads, eluted using 2x SDS loading buffer) were separated on a 10% SDS gel that was Coomassie blue stained afterwards. Bands were excised and further processed for mass-spectrometry analysis. **F**. Identification of MK2 peptides from mass-spectrometry (1E.). The three translation initiation sites are marked in bold and italics. Red sequences depict detected peptides and black ones the non-detected.

To test the hypothesis of the existence of alternative translation products, we exploited existing RNA sequencing (RNASeq) and ribosome footprinting (RiboSeq) data from different cell lines. Initially, our own data from murine bone marrow-derived macrophages (BMDMs) (23) were reanalyzed and revealed the existence of an MK2 transcript with an extended 5′UTR that also contained several ribosome footprints (RFPs) within this region (Fig. 1C). This supported the idea of the existence of alternative translation events in the 5′UTR. In fact, an extended MK2 transcript that corresponded to the transcript length we detected by RNASeq, was recently deposited at NCBI (NM_008551.2) and replaced the previously annotated transcript having only a very short 18 nt 5′UTR (NM_008551.1; Fig. 1C). The analysis of this transcript showed a potential in-frame extension of the hitherto known MK2 ORF in the upstream direction (Fig. 1D). Within this extension we found two putative alternative translation initiation start sites (aTIS) at positions –141 and –309, coding for Leucine (CUG) and Valine (GUG), respectively. The CUG and GUG are surrounded by the alternative Kozak-sequences GCCGAGC*<u>CUG</u>*GAG and GCAGCCT*<u>GUG</u>*ACG (3).

Consistent with the Western blot results, RFPs and sequences analyses revealed an in-frame extension in the 5′UTR of the human MK2 transcript bearing the two alternative start codons CUG and GUG (Fig. S2A,B). Interestingly, in the human transcript, the CUG is in almost the same distance to the AUG (-144bp, –48Aa). Overall, more than 80% sequence identity within the first 200bp upstream of the start AUG codon exists between the murine and human 5′UTR (Fig. S2C). Further sequence analyses showed that the 5′UTR extension is evolutionary conserved among species, indicating a distant origin for alternative translation of MK2 isoforms (Supplementary File S1). This observation was supported by analyzing published RiboSeq data sets of murine and human cells that were either treated with cycloheximide, harringtonine or lactimidomycin before sequencing of ribosome-protected mRNA fragments (8,24–29). Since mRNA bound ribosomes are frozen at translation initiation start sites by these treatments as previously exemplified in pioneering RiboSeq studies (30,31), the existence of footprints in the 5′UTR of MK2 strongly argues for alternative translation events.

Indeed, we could observe such footprints in murine and human cell lines in the 5′UTR of MK2 (Fig. S3 and 4).

To validate the existence of an N-terminally extended MK2 isoform due to alternative translation initiation, we sought to purify endogenous MK2 from murine RAW264.7 cells that express almost equal levels of both MK2 isoforms (Fig. 1A,E). By using a MK2-Nanotrap (Chromotek, München, Germany) we were able to enrich and purify considerable amounts of endogenous MK2 from these cells and subsequently analyzed the purified protein by mass-spectrometry (Fig. 1E,F). We readily identified peptides derived from the predicted in-frame 5′UTR extension originating from the CUG aTIS at position –141 (Fig. 1F). The detection of peptides derived from sequences further upstream of this CUG indicates that also the GUG in position –309 or other codons may be used for alternative translation initiation. In sum, these results confirmed the existence of at least one aTIS at position –141 in the 5′UTR of the MK2 transcript resulting in the translation of a minimum of two distinct MK2 isoforms in murine and human cells from one mRNA.

**Figure 4.**
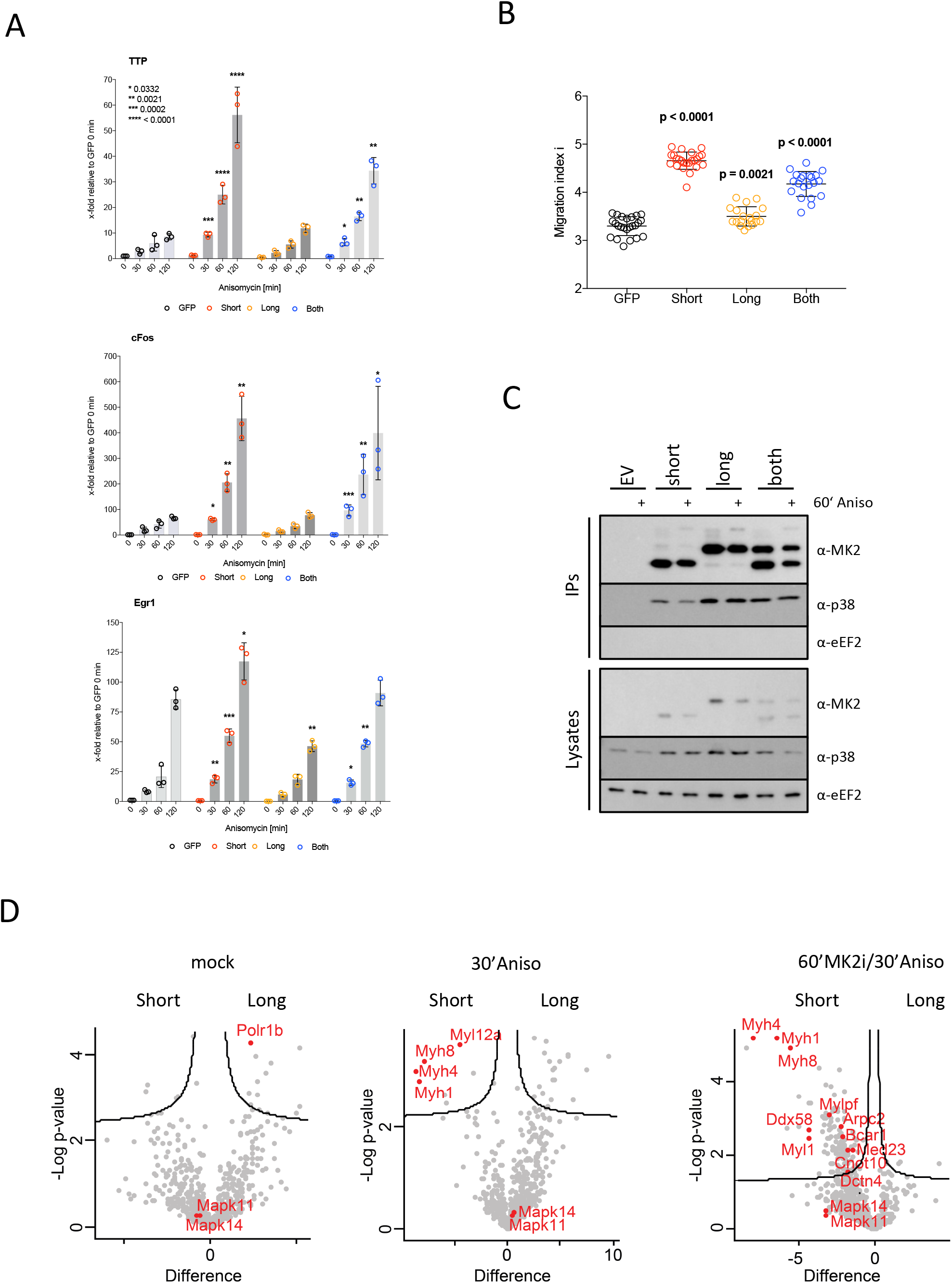
Phenotypical consequences of MK2 isoform expression. **A**. Analysis of TTP, cFos and Egr1 expression upon Anisomycin stimulation for the indicated times in differentially rescued MK2-deficient MEFs. Each data point represents a biological replicate that was measured in duplicates. Error bars represent the mean of three independent experiments (n = 3) +/**−** the standard deviation. P-values were retrieved from one-way ANOVA testing using the PRISM 7 software and are related to GFP/EV-rescued MEFs. **B**. Fluorescence-based scratch wound healing assay for differentially rescued MK2-deficient MEFs as shown before (34). Each individual data point represents a biological replicate (n > 18) of the analysis and the mean of all replicates is shown with its standard deviations. P-values were retrieved from one-way ANOVA testing using the PRISM 7 software and are related to GFP/EV-transduced MK2-deficient MEFs. **C**. The interaction of MK2 isoforms with their endogenous upstream activating kinase p38^MAPK^ was detected by co-immunoprecipitation experiments. Non-stimulated and Anisomycin-stimulated MK2/3 DKO complemented MEFs were analyzed. A MK2-Trap was used to precipitate MK2 effectively. Beads were eluted using 2x SDS loading buffer and samples were analyzed by Western Blot development for MK2 and p38^MAPK^. eEF2 development served as a control. **D**. Volcano plots of significantly MK2 short and MK2 long interacting proteins at the indicated conditions. Selected proteins are highlighted in red. The black lines define the border of significantly and non-significantly interacting proteins.

### Mechanistic insights into regulation of MK2 translation

To further characterize the usage of the putative aTIS at position –141 of MK2 mRNA, we fused the extended endogenous murine 5′UTR with the known ORF sequence of murine MK2 and expressed the construct in HeLa and U2OS cells. The high GC-content of the 5′UTR (>85%) prompted us to introduce a synthetic DNA flanked by restriction sites (Fig. S5A). Expression of the 5′UTR-MK2-ORF construct resulted in the two-band pattern seen for endogenous murine MK2 independent of the expressing cell line (Fig. 2A, S5B). Mutagenesis of the putative aTIS in positions –141 and –309 resulted in an altered band pattern and revealed that the triplet CUG at –141 is used for alternative translation initiation under these conditions leading to expression of the long isoform (Fig. 2A). Of note, destroying the alternative Kozak-sequence surrounding this aTIS (mutation GAG to CAU) impaired the expression of the long isoform (Fig. 2A, lane 6). Interestingly, it was even possible to generate an extralong MK2 isoform (super-long) by mutating –309 GUG to AUG (Fig. 2A, lane 7).

**Figure 2.**
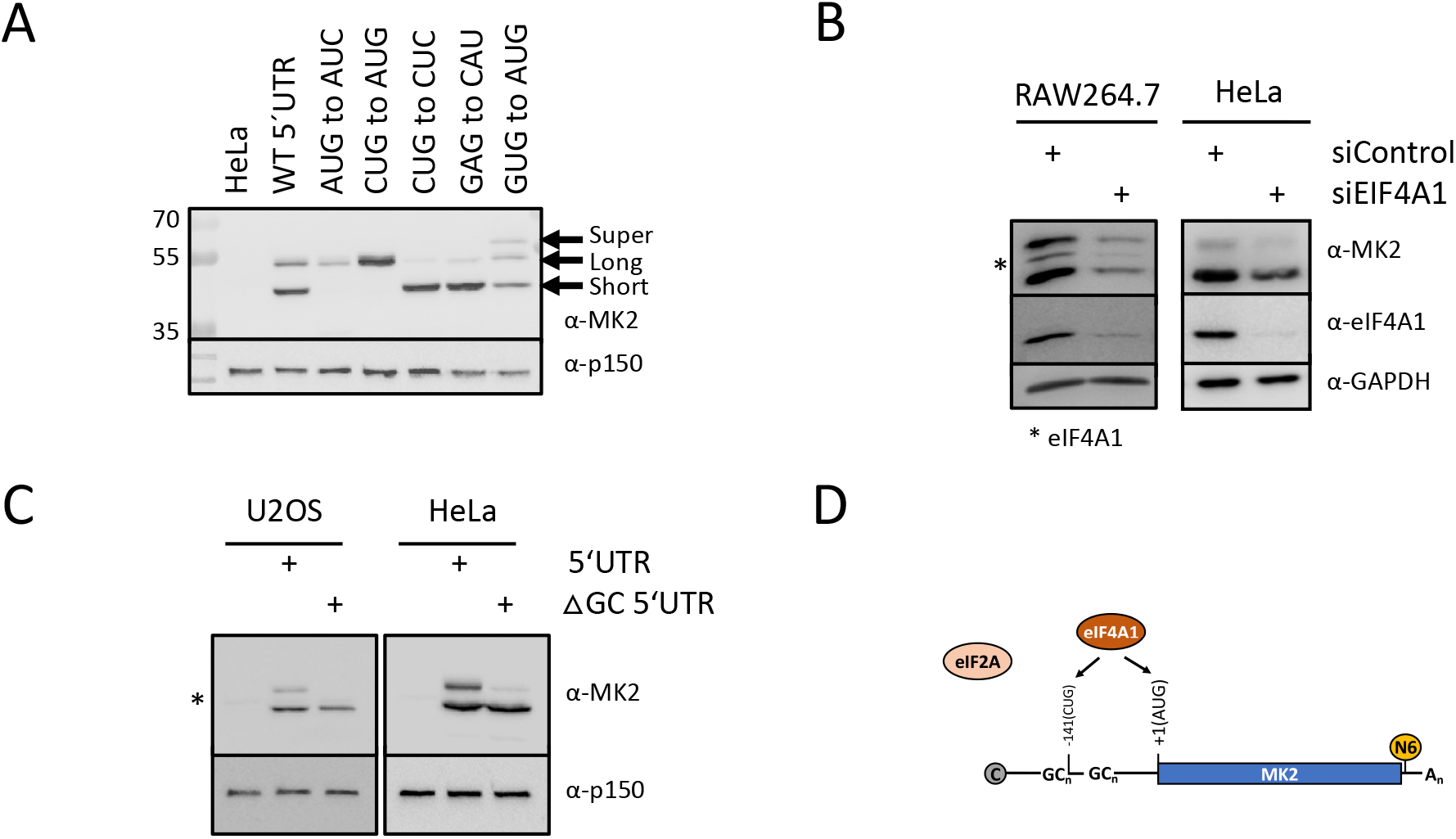
Mechanistic insights into regulation of MK2 translation. **A**. Expression of murine MK2 5′UTR isoforms in HeLa cells through introduction of point mutations. After cloning the entire 5′UTR of murine MK2 into peGFP-C1 based vector, different point mutations were introduced into the 5′UTR (see Figure S5). HeLa cells were transfected with these plasmids and lysates were analyzed by Western Blot with an anti-MK2 antibody and an anti-p150 antibody to control equal loading. Arrows indicate for the different MK2 isoforms resulting from point mutations within the 5′UTR. **B**. RAW264.7 and HeLa cells were transfected with an siRNA targeting eIF4A1. Lysates were then analyzed by Western Blot against eIF4A1 to monitor knockdown efficacy and MK2 to detect the endogenous levels of MK2. GAPDH as loading control was used to normalize the signals for quantification (see graph next Western Blot panels). The * indicated for the eIF4A1 band. **C**. Translation of murine MK2 isoforms upon deletion GC-rich sequences upstream of the CUG aTIS in HeLa and U2OS cells. HeLa and U2OS were either mock-transfected or transfected with the constructs represented next to the Western Blot panels. Lysates were analyzed by Western Blot using an anti-MK2 antibody and an anto-p150 as a loading control. The * indicates for the weak band of the short endogenous human MK2 isoform. **D**. Schematic representation of the factors that can influence MK2 translation. We confirmed the need of eIF4A1 to promote translation from the CUG aTIS in the 5′UTR of MK2. Adenosine Methylation near the stop codon, as shown indirectly by the knockdown of the methyltransferase Mettl3, and depletion of eIF2A had no effect on MK2 translation in our hands.

Next, we explored the underlying molecular mechanism responsible for differential MK2 isoform translation *in vivo*. Human cells treated with the anti-cancer drug silvestrol respond with a decrease in the translation of a selective set of mRNAs due to inhibition of the RNA helicase eIF4A1 (32,33). Amongst them, MK2 mRNA is a major target (33). Additionally, it was shown that mRNAs commonly regulated by this mechanism, contain long and GC-rich 5′UTRs with G-rich quadruplex structures (32,33). We were able to detect putative sequences that give rise to such structures within the 5′UTR of human and murine MK2, located within the coding and assumed non-coding region of the elongated 5′UTR (Table S1). We therefore speculated that the helicase eIF4A1 might play a role in selective MK2 isoform translation. Upon depletion of eIF4A1 in murine and human cells a marked decrease in translation of both MK2 isoforms was observed (Fig. 2B, S6A). Similar results were obtained in WT MEFs, NIH3T3 and HEK293T cells (data not shown). To additionally prove a role for the GC-rich clusters in isoform translation, we generated a mutant of the above-mentioned construct lacking the first three GC-rich clusters (positions 21, 96 and 128, see Table S1 and schematic representation of the constructs, Fig. S6B) upstream of CUG in –141 position (DGC 5′UTR). These deletions were accompanied by the almost complete abrogation of long isoform translation (Fig 2C). It was proposed that the translation initiation auxiliary factor eIF2A is important for non-AUG translation initiation (7) and expression of PTEN-alpha isoforms are dependent of this factor (34,35). However, we did not observe any effect of eIF2A depletion on MK2 isoform translation (Fig. S6C). Epigenetic markers of mRNAs, such as m^6^-adenosine methylation, influence stability and translation of modified mRNAs (36). Human MK2 mRNA is heavily methylated near the stop codon and translation of MK2 mRNA was reported to be strongly decreased upon depletion of the methyltransferase Mettl3 (37,38). However, we were not able to support these findings, as knockdown of Mettl3 in murine and human cell lines did not change the normal pattern of MK2 translation (Fig. S6D). In summary, we demonstrate a role for the RNA helicase eIF4A1 and putative G-rich quadruplex structures in the generation of distinct MK2 isoforms from a single mRNA species (cf. Fig. 2D).

### Functional characterization of MK2 isoforms

In order to address the physiological consequences of MK2 isoform expression, we stably reconstituted MK2-deficient mouse embryonic fibroblasts (MEFs) cell lines (13) with the short, the long, the super-long isoform or with both (long and short) isoforms of MK2 (Fig. 3A). First, we assessed the relative stability of the two MK2 isoforms after a cycloheximide-induced block of *de novo* protein translation (Fig. 3B). As seen, the longer variant was considerably more stable than the short isoform in this assay. Both GFP-tagged isoforms localized similarly in non-stressed cells (Fig. S7A), and when we activated p38^MAPK^ by anisomycin stress, we observed their similar redistribution to the cytosol confirming previous observations (summarized in (39)). Monitoring the redistribution of the super-long isoform was not possible since transfection of this construct was accompanied by short and long isoform expression (Fig. S7A). We then tested the ability of the isoforms to phosphorylate the known MK2 substrates Reticulon 4B/NOGO-B and Hsp27 (also HSBP1 or Hsp25 in mice) upon activation by anisomycin in cells (Fig. 3C, S7B). The specificity of phosphorylation was assessed with chemical inhibitors of p38^MAPK^- and MK2 (Fig. 3C). While NOGO-B was readily phosphorylated by both isoforms, phosphorylation of the small heat shock protein Hsp27, a bona fide substrate of MK2, could only be detected for the short MK2 isoform, (Fig. 3C). This finding indicated the intriguing possibility that both isoforms differed in their substrate specificity.

**Figure 3.**
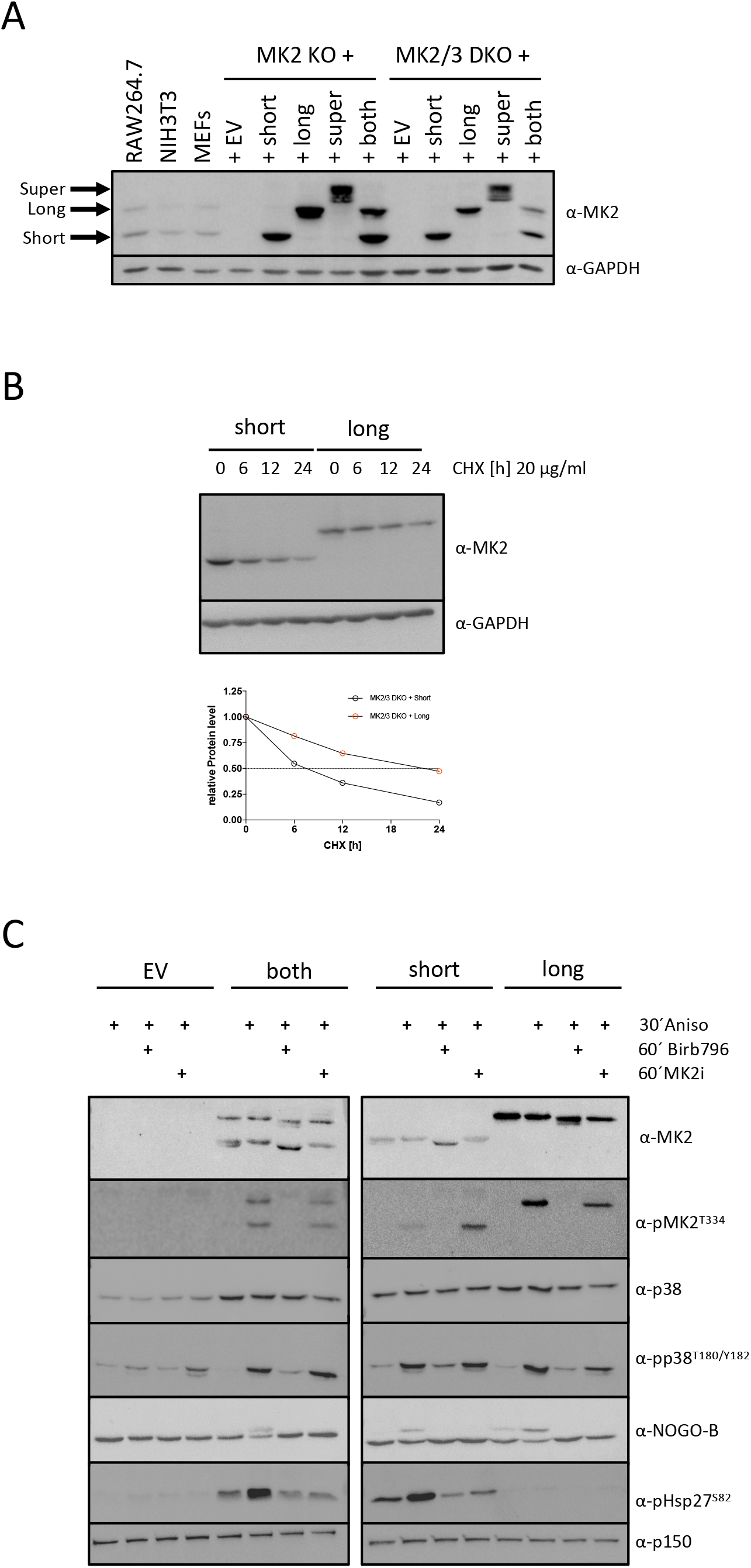
Functional characterization of MK2 isoforms. **A**. Complementation of MK2 KO and MK2/3 DO mouse embryonic fibroblasts (MEFs) by retroviral transduction. Subcloning of 5′UTR mutants into pMMP-MK2-IRES-eGFP based vectors (13) and subsequent viral transduction of MK2-deficient and MK2/3 double-deficient MEFs resulted in expression of either the short or the long MK2 isoform only, both isoforms in parallel (both) or a super-long (super) isoform. Empty vector transduced cells served as controls and non-transduced murine cell lines (RAW264.7, NIH3T3 and wildtype (WT) MEFs) indicated the position of endogenous MK2 isoforms and expression levels in transduced cell lines. **B**. The stability of MK2 isoforms in MK2 KO or MK2/3 DKO complemented MEFs was analyzed after incubating cells with 20 µg/ml cycloheximide (CHX) for the indicated times. A quantification of the band intensities can be found in Figure S6A. The GAPDH development served as a loading control. **C**. The differentially complemented MK2/3 DKO MEFs from Fig. 3A were analyzed for the phosphorylation of known MK2 substrates (NOGO-B and Hsp27) upon Anisomycin stimulation. Phosphorylation of the MK2 upstream p38^MAPK^(pp38^T180/Y182^) and MK2 (pMK2^T^334) itself was used as a readout for activation of the pathway. To control activity, cells were either pre-incubated with an p38^MAPK^ inhibitor (Birb796) or a MK2 inhibitor (PF-3644022). p150 development served as loading control.

Taken together we were able to reconstitute MK2-deficient cells with individual isoforms. Both murine versions of MK2 share molecular properties such as the ability to translocate from the nucleus to the cytoplasm but differ in their stability and substrate specificity.

### Phenotypical consequences of MK2 isoform expression

Expression of both isoforms differ markedly among mouse tissues (Fig. 1A,B). Therefore, we speculated on physiological relevant differences and attempted to explore the phenotypical consequences of isoform expression in MK2- and MK2/3-deficient MEFs. We initially tested proliferation of the differently rescued cell lines (Fig. S8). Cells grew to a similar extent independently of the isoform used for rescue. We then checked the induction of the IEGs early growth response 1 (Egr1), FBJ osteosarcoma oncogene (cFos) and tristetraprolin (TTP or Zfp36) upon anisomycin stimulation of MK2/3-deficient MEFs rescued with the different isoforms of MK2 (Fig. 4A). While expression of the short isoform markedly elevated the expression of all of these genes, the long isoform of MK2 was not able to induce IEGs. Of note, we were not able to detect differences in SRF phosphorylation by the isoforms (data not shown) indicating that other MK2-dependent mechanisms regulate IEG transcription under these conditions. Since it is known that the N-terminus of MK2 plays an important role in cell motility (40), we also assayed for cell migration in the rescue cell lines using a scratch assay (Fig. 4B). As seen, the MK2-associated migration phenotype could only be rescued by the short but not the long MK2 isoform, consistent with our observation that only the short isoform of MK2 is competent for phosphorylation of the migration associated substrate Hsp27 (Fig. 3C).

Collectively these results highlighted differential functional properties of alternatively translated MK2 isoforms.

### MK2 isoform interaction profiling

The phenotypic differences in our rescue cell lines suggested that the two MK2 isoforms could have overlapping as well as unique interaction partners. As a control for the proficiency of both proteins, we first immunoprecipitated the isoforms with MK2-Trap beads and confirmed their well-characterized interaction with the upstream activating kinase p38^MAPK^ (Fig. 4C). We then used quantitative mass-spectrometry to analyze the interaction partners of both isoforms in cells that were left untreated or treated with combinations of anisomycin and MK2 inhibitor (MK2i: PF-3644022). This approach allowed us to compare MK2 activity-dependent interactions partners of the two MK2 isoforms. While we detected only few isoform-specific interactions in resting cells, stress-induction by anisomycin, especially in the combination with MK2 inhibitor, revealed a strong bias towards the short MK2 isoform for interaction with proteins of the myosin family (Myh1, Myh4, Myh8, Myl1, Myl12a and Mylpf) and the transcriptional machinery (Med23, Cnot10) (Fig. 4D, Supplementary File S2). GO term analysis (data not shown) revealed more proteins, such as Smtn, Arpc2 or Ddx58, involved in migration to selectively interact with the short MK2 isoform (Fig. 4D). Given the fact that MK2-dependent migration is only rescued by the short isoform, which is also uniquely capable of phosphorylating Hsp27 upon stress (Fig. 3B), we speculate that these short isoform-specific interactions underlie the migration of MK2-proficient cells. In line with our biochemical experiment (Fig. 3C), our proteomics analysis confirmed that both MK2 isoforms readily interact with p38alpha/beta (Mapk11/14) in a stimulus-independent manner (Fig. 4D).

Taken together, these results revealed that the two MK2 isoforms have both common and distinct interaction partners. The specific interactions of the short isoform with migration-associated proteins and transcriptional regulators may explain why some MK2-dependent phenotypes are only rescued by this MK2 variant.

## Discussion

The p38^MAPK^-activated kinase MK2 is a central regulator of immediate early genes transcription and post-transcriptional gene regulation. Here we show through a combination of *in silico* and experimental methods that murine and human MK2 itself is regulated post-transcriptionally at the level of translation. A long isoform of the protein is constitutively and alternatively translated from an in-frame CUG initiation site (aTIS) upstream of the known AUG initiation codon that gives rise to the known canonical short isoform of MK2. The conserved 5′UTR of the MK2 transcript has a high GC-content and the usage of the CUG aTIS is potentially dependent on surrounding G-rich quadruplex structures. Consequently, translation of both isoforms is dependent on the RNA helicase eIF4A1.

By introducing point mutations in either the CUG or the AUG we were able to change the preference of translation initiation site. The aTIS CUG in position –141 is surrounded by an alternative Kozak-sequences (3). Importantly, translation starting from this CUG was abrogated when the adjacent downstream GAG triplet was mutated into CAU, confirming the need for the Kozak-sequence. Our initial *in silico* analyses of ribosome footprinting data prompted us to investigate alternative translation in the previously uncharacterized 5′UTR of the murine MK2 transcript. We identified ribosome footprints around the CUG triplet in the human and murine MK2 5′UTR in cells treated either with cycloheximide, lactimidomycin and harringtonine. This showed us that we not only detect translating, but also initiating ribosomes occupying these sites. Detection of endogenous peptides by mass-spectrometry that could be mapped to the potential in-frame sequence upstream of the AUG start codon supported the *in silico* analyses.

Two reports identified MK2 as a target of eIF4A1 with G-rich quadruplex structures in its 5′UTR, by either knocking down or inhibiting the RNA helicase with the anticancer drug silvestrol (32,33). RNA G-rich quadruplex structures in 5′UTRs are known to influence translation (41). We found six potential sites within the full and extended 5′UTR (Table S1). The human MK2 sequence bears the optimal 12-mer G-rich quadruplex sequence and its translation efficiency is strongly decreased upon treatment with silvestrol (33). Indeed, our data confirms that eIF4A1 and GC-rich sequences that might form G-rich quadruplex structures regulate MK2 translation. More specifically, these sequences upstream of the AUG translation initiation site, lead to the usage of a CUG triplet as an alternative translation initiation site within the 5′UTR of the MK2 mRNA. By deleting the GC-rich sequences upstream of the CUG, translation of the high molecular weight MK2 isoform was abrogated. The data could suggest that the scanning mechanism of the ribosome is slowed down and consequently a leaky scanning leads to the usage of the CUG for translation initiation. This mechanism appears evolutionarily conserved, yet murine cells produce relatively more of the long and alternative isoform compared to their human counterparts. This correlates with the fact that the human 5′UTR is shorter than in the murine transcript and the CUG is only surrounded by one putative upstream G-rich quadruplex structure. The resulting decrease in structural barriers is likely accompanied by less translation from the aTIS and, consequently, the production of relatively lower amounts of the long MK2 isoform in human cells. Of note, the balance between MK2 isoform production differs markedly between murine tissues and cell lines, suggesting that these proteins have some non-overlapping functions *in vivo*. mRNA adenosine methylation (N6-methyladenosine or m^6^-A, respectively) of the MK2 mRNA near the stop codon by METTL3 was reported to lower translation without affecting the mRNA level, and this effect was independent of m^6^-A reader proteins such as YTHDF1,2 and 3 (37,38). In the present study, we could not confirm an involvement of METTL3 in general or specific MK2 isoform translation. Examples of in-frame N-terminal protein extensions originating from alternative translation starting in the 5′UTR have previously been described i.e. for PTEN (34,35) or Thioredoxin/Glutathion (42). It was suggested that eIF2A is a specificity factor for the translation of the long PTEN alpha isoform (34,35) and a general factor for non-AUG translation initiation (7,8). To this end we could not observe an involvement of eIF2A in elongated MK2 isoform translation. The model in Figure 2D summarizes our investigations on MK2 translation and it remains to be clarified whether the RNA helicase Ddx3 (6) is a regulatory factor for MK2 isoform translation alone or in cooperation with eIF4A (43).

The two MK2 isoforms identified in this report are highly similar in terms of their activation by and interaction with the upstream kinase p38^MAPK^. The long MK2 isoforms had a decreased turnover as compared to its short counterpart, but it failed to phosphorylate the MK2 *bona fide* substrate Hsp27 upon anisomycin stimulation. This is similar to the different p38^MAPK^ isoforms, that exhibit distinct substrate specificities although displaying a high degree of amino acid sequence identity (44). Testing potential phenotypical consequences of selective isoform expression in an MK2-deficient background revealed that only reintroduction of the short isoform or both isoforms in parallel (equal to the wildtype situation) can rescue migration defects and IEGs expression. By determining the interaction partners of MK2 isoforms, we were able to shed some light on these phenotypical differences. The preferred binding of the short MK2 isoform to proteins of the cytoskeleton and the transcription machinery correlates with the unique ability of this isoform to rescue migration and transcriptional defects. Both processes are well-established MK2-dependent responses (15,45) and the importance of the canonical MK2 N-terminus was confirmed here (40). Although not elucidated in this study, the long isoform of MK2 is likely to have unique functions as well, which are not supported by the canonical isoform. Of note, the unique N-terminal extension of this protein contains putative proline-directed serine and threonine phosphorylation sites, that we could detect in their phosphorylated form in our mass-spectrometry data set. Interestingly, the phosphorylation of these sites seems not induced by anisomycin-stimulated p38^MAPK^ activation. Such additional and isoform-specific modifications could potentially allow for the recruitment of unique substrates, and the assignment of unique functions of the long MK2 isoform clearly warrants future investigation. We therefore entertain the possibility that the extended N-terminus is targeted by other proline-directed kinases than p38^MAPK^ such as cyclin-dependent kinases (CDKs) (46). We can only speculate on the consequences of extended N-terminus phosphorylation, but want to stress that expression of the short isoform alone was often accompanied by a “super-rescue” of MK2-deficiency compared to the rescue by expression of both isoforms together (see Fig. 4A,B).

Future phospho-proteomic studies could shed additional light on substrate specificities for the two MK2 isoforms and their consequences *in vivo*. In addition, the generation of gene-edited mice that exclusively express either MK2 isoform could help to resolve these outstanding issues.

In conclusion, our work reveals the abundant alternative translation of a hitherto unrecognized MK2 isoform with distinct functional properties, highlighting that the biology of this important kinase is still not fully understood. Our work also resolves a long-standing mystery surrounding the presence of the additional high-molecular weight MK2 band often detected in Western blot (13,21,22).

## Experimental Procedures

### Tissue culture, cell lines, treatments

Cells were grown on plates in DMEM or in RPMI for suspension cells in the presence of 10 % (v/v) FCS, 100 U penicillin G/mL, and 100 mg/mL streptomycin (all purchased from Life Technologies) in a humidified atmosphere at 37°C with 5% CO_2_ supplementation. Treatments with stimulators and inhibitors were done at the following final concentrations: Anisomycin, 10 µg/ml, dissolved in DMSO, Sigma; BIRB796, 1 µM, dissolved in DMSO (Axon Medchem); MK2 inhibitor PF3644022 (10µM, Sigma) dissolved in DMSO; Cycloheximide, 20 µg/ml, dissolved in water, Sigma. For gene expression experiments cells were starved in serum-free medium for a minimum of 8h. MK2-deficient and MK2/3 double-deficient MEFs were described before (13,15). All other standard laboratory cell lines were either kind gifts or obtained from ATCC.

### Isolation of mouse organs/bone marrow-derived macrophages

Organs of a wildtype and bone-marrow of wildtype, MK2 KO and MK2/3 DKO animals (C57BL/6 background) (12) were isolated after sacrificing them. Handling of mice complied with all relevant ethical regulations and was approved by the ethics committees of Hannover Medical School. Organs were flash-frozen in liquid nitrogen before lysis. To derive macrophages from bone marrow after the femurs of one mouse of each genotype were flushed. Cells were then grown in the presence of 10 ng/ml M-CSF for approximately 9–10 days before using them for further experiments (14).

### Transfection of cell lines

Cells were seeded the day before transfection to reach 50% density on the next day. The medium was then changed to penicillin/streptomycin-free. DNA was mixed in Opti-MEM (Life Technologies) and complexed with polyethylenimine (PEI, 1 µg/ml, pH 7.0, Sigma). After a minimal incubation of 10 min at room temperature the mix was added drop-wise to the cells. Medium was changed to full medium 6 h and cells were treated 24 h posttransfection and harvested.

For microscopy experiments using U2OS cells transfections were carried out according to the manufacturer using FUGene6 (Promega).

### siRNA transfections

All cell lines were transfected using HiPerFect (Qiagen), RNAiMAX (Thermo Fisher Scientific) or Lipofectamin 2000 (Thermo Fisher Scientific/Invitrogen) according to the manufacturer’s instruction. The following siRNAs were used:

AllStars Negative Control siRNA, Qiagen SI03650318, murine sieIF4A1: SMARTpool siGENOME (M-060466–01-0005), Dharmacon, human siEIF4A1:

CUGGCCGUGUGUUUGAUAUdTdT (47), murine siMettl3 #1:

AGCUACCGUAUGGGACAUUAA (Qiagen FlexiTube siRNA SI01304135 Mm_Mettl3_1), murine siMettl3 #2: CUGGACUGCGAUGUGAUUGUA (Qiagen FlexiTube siRNA SI01304142 Mm_Mettl3_2), human siMETTL3 #1: CAGGAGAUCCUAGAGCUAUUA (Qiagen FlexiTube siRNA SI04241265 Hs_METTL3_6), human siMETTL3 #2:

CUGCAAGUAUGUUCACUAUGA (Qiagen FlexiTube siRNA SI04317096 Hs_METTL3_7), murine sieIF2A #1: UGGAAAUCUUCGAGGACAAdTdT, murine sieIF2A #2: UGAAUUUGUUGGCAGGAAAdTdT, human siEF2A #1: AGAAAGUGCUGUAGUGCAAdTdT, human siEIF2A #2:

UGUUAAACCCAUUGGUAUAdTdT. siRNAs were either synthesized by Eurofins/MWG or purchased from the indicated supplier.

### Generation of stable shRNA knockdown cell lines

Stable HT29 cells were previously described (48). Two different lentiviral particles (Mission shRNA lentiviral particles (Sigma), #TRCN0000002282 and # TRCN0000002283) together and a control lentiviral particle (Non-Target shRNA Control) were used. After transduction, cells were selected with puromycin (2 µg/mL) for 2 weeks and consequently tested for MK2 expression by Western Blot.

### Cloning

A detailed description of the cloning strategy to express MK2 with its full-length 5′UTR is shown in Fig. S5A. The indicated 441 bp 5′UTR-MK2 fragment having NheI and PstI sites was synthesized by GENEART GmbH (Thermo Fisher Scientific). The peGFP-C1-MK2 and pMMP-MK2-IRES-eGFP constructs for expression in mammalian cells and for retroviral transduction were described before (13,49).

### SDS-PAGE and Western Blot

If not stated differently proteins were separated on 10% SDS acrylamide gels and transferred to nitrocellulose membrane by semi-dry transfer. For Western Blot developments blocking and secondary antibody incubations were done for 1h at room temperature in 5% dry-milk powder dissolved in PBS containing 0.1% Tween 20 (PBS/T). Primary antibodies were usually incubated overnight at 4°C and diluted in 5% BSA dissolved in PBS/T. Antibodies to detect specific proteins: anti-MK2 (CST, #3042 and #12155), anti-pMK2^T222^ (CST, #3316), anti-pMK2^T334^ (CST, #3041), anti-MK3 (CST, #3034), anti-p38 (CST, #9012), anti-pp38^T180/Y182^ (CST, #4511), anti-NOGO-B (R+D Systems, AF6034), anti-Hsp27^S82^ (CST, #9709), anti-GAPDH (EMD Millipore, #MAB374), anti-Tubulin (Sigma, #T6199), anti-eEF2 (SCBT, sc-13004), anti-p150 (BD Biosciences, #610473), anti-Mettl3 (CST, #96391), anti-eIF4A1 (CST, #2490T), anti-eIF2A (Proteintech, 11233–1-AP), anti-GFP (SCBT, sc-9996) and anti-TTP (N-Term) (Sigma T5327) HRP-coupled secondary antibodies were used for ECL-based detection of proteins on either a LAS3000 FujiFilm or a Bio-Rad ChemiDoc Imaging System.

### MK2 precipitation and LC-MS for the identification of MK2 peptides

Precipitation MK2 was done with the help of an agarose MK2-Trap (Chromotek) following the manufacturer’s protocol. After final washing of the agarose beads, proteins were eluted using 2x SDS loading buffer and incubated for 5 min at 95°C. Eluted proteins were then alkylated by addition of acrylamide up to a concentration of 2% and incubation at room temperature for 30 min. Samples were then separated on a 10% SDS acrylamide gel. After electrophoresis proteins were stained with Coomassie Brilliant Blue (CBB) for 15 min and background staining was reduced with water. Specific MK2 lanes were excised and minced to 1 mm³ gel pieces. Further sample processing was done as described (50). Briefly, gel pieces were destained two times with 200 µL 50% acetonitrile (ACN), 50 mM ammonium bicarbonate (ABC) at 37°C for 30 min and then dehydrated with 100% ACN. Solvent was removed in a vacuum centrifuge and 100 µL 10 ng/µL sequencing grade Trypsin (Promega) in 10% ACN, 40 mM ABC were added. Gels were rehydrated in trypsin solution for 1 hour on ice and then covered with 10% ACN, 40 mM ABC. Digestion was performed overnight at 37°C and was stopped by adding 100 µL of 50% ACN, 0,1% trilfluoroacetic acid (TFA). After incubation at 37°C for 1 hour the solution was transferred to a fresh sample vial. This step was repeated twice and extracts were combined and dried in a vacuum centrifuge. Dried peptide extracts were redissolved in 30 µL 2% ACN, 0.1% TFA with shaking at 800 rpm for 20 min. After centrifugation at 20,000 x g aliquots of 12.5 µL each were stored at –20°C. Peptide samples were separated with a nano-flow ultra-high pressure liquid chromatography system (RSLC, Thermo Scientific) equipped with a trapping column (3 µm C18 particle, 2 cm length, 75 µm ID, Acclaim PepMap, Thermo Scientific) and a 50 cm long separation column (2 µm C18 particle, 75 µm ID, Acclaim PepMap, Thermo Scientific). Peptide mixtures were injected, enriched and desalted on the trapping column at a flow rate of 6 µL/min with 0.1% TFA for 5 min. The trapping column was switched online with the separating column and peptides were eluted with a multi-step binary gradient: linear gradient of buffer B (80% ACN, 0.1% formic acid) in buffer A (0.1% formic acid) from 4% to 25% in 30 min, 25% to 50% in 10 min, 50% to 90% in 5 min and 10 min at 90% B. The column was reconditioned to 4% B in 15 min. The flow rate was 250 nL/min and the column temperature was set to 45°C. The RSLC system was coupled online via a Nano Spray Flex Ion Soure II (Thermo Fisher Scientific) to an LTQ-Orbitrap Velos mass spectrometer. Metal-coated fused-silica emitters (SilicaTip, 10 µm i.d., New Objectives) and a voltage of 1.3 kV were used for the electrospray. Overview scans were acquired at a resolution of 60k in a mass range of m/z 300–1600 in the orbitrap analyzer and stored in profile mode. The top 10 most intensive ions of charges two or three and a minimum intensity of 2000 counts were selected for CID fragmentation with a normalized collision energy of 38.0, an activation time of 10 ms and an activation Q of 0.250 in the LTQ. Fragment ion mass spectra were recorded in the LTQ at normal scan rate and stored as centroid m/z value and intensity pairs. Active exclusion was activated so that ions fragmented once were excluded from further fragmentation for 70 s within a mass window of 10 ppm of the specific m/z value. Raw data were processed using Proteome discoverer (Thermo Fisher Scientific) and murine uniprot/swissprot data bases containing common contaminants. Proteins were stated identified by a false discovery rate of 0.01 on protein and peptide level and quantified by extracted ion chromatograms of all peptides.

### MK2 interaction profiling by label-free mass-spectrometry

Per sample 3 to 4 confluent 15 cm dishes of the corresponding MK2 MEFs were either mock-treated (DMSO), pre-treated with the MK2 inhibitor PF3644022 (10µM, Sigma) for 60 minutes or stimulated with Anisomycin (1µg/ml, Sigma) for 30 minutes before lysis. Approximately 5 mg of lysate was administered to MK2-immunoprecpitations using an agarose MK2-Trap binding resin (Chromotek). Beads were eluted in 2x Laemmli buffer and the resulting supernatants were loaded onto a 4–12% NuPAGE Bis-Tris Gel (Novex, Thermo Fisher Scientific). Gels were then fixed and stained using the colloidal blue staining kit (Life Technologies/Invitrogen). Gel lanes were sliced into five pieces and proteins were digested in-gel using standard methods (51). Peptides were reconstituted in 5% CAN 0.1% TFA and analyzed with an Easy-nLC 1200 system (Thermo Scientific) connected to a Fusion Lumos Orbitrap mass spectrometer (Thermo Scientific) through a nanoelectrospray ion source. Peptides were separated on a 15 cm (75 µm inner diameter) in-house column packed with 1.9 µm reversed-phase C18 beads (Reprosil-Pur AQ, Dr. Maisch), with 45 min gradients. Spray voltage was set to 2kV, capillary temperature to 275°C and RF level to 30%. A full scan was performed at a resolution of 60,000, with a scan range of 350–1400 m/z, a maximum injection time of 20 ms and with an AGC target of 500,000. Precursors were isolated with a width of 1.3 m/z and precursor fragmentation was accomplished using higher collision dissociation (HCD). Only precursors with charge state 2–5 were considered. MS/MS spectra were acquired in the Orbitrap, and settings included a loop count of 7, a maximum injection time of 118 ms and a resolution of 60,000. Raw MS data were analyzed using MaxQuant software (version 1.6.0.17). Proteins were identified with parameters previously described (52) and quantified using the LFQ algorithm integrated in MaxQuant (53). Match between runs function was used. Data analysis was performed using Perseus software (54). Missing values were imputed in all samples using Perseus default settings. Volcano plots were generated by plotting the – log10 transformed FDR-adjusted p-values derived from a two-sided t-test versus log2 transformed fold changed. Significance was determined based on a hyperbolic curve threshold of S0=0.1 using Perseus.

### Migration Assay

Fibroblast migration was analyzed by a fluorescence-based scratch wound-healing assay as described previously (45).

### WST-1 Assay

Cell viability assays using WST-1 (water soluble tetrazolium, Roche # 11 644 807 001) were performed as described before (14).

### RNA extraction and qPCR

Total RNA from stimulated cells was extracted using Trizol (Thermo Fisher Scientific/Invitrogen) according to the instructions. 500 ng RNA were then reverse transcribed to cDNA using random hexameric primers and Reverse Transcriptase (Thermo Fisher Scientific) in the presence of RNase inhibitors. Quantitative PCRs (qPCRs) were performed on One Step-Plus real-time PCR systems (Applied Biosystems/Thermo Fisher Scientific) using a 2x SYBR Green no ROX qPCR mix (Bioline) in a two-step cycling program. The following specific qPCR primer were used to detect mRNA level: cFos-fwd: CTACTGTGTTCCTGGCAATAG, cFos-rev: AACATTGACGCTGAAGGACTA, Egr1-fwd: ACAGAAGGACAAGAAAGCAGAC, Egr1-rev: CCAGGAGAGGAGTAGGAAGTG, TTP-fwd: CATCTACGAGAGCCTCCAGTC, TTP-rev: ACGGGATGGAGTCCGAGTTTA. Specific signals were normalized to GAPDH or beta-Actin mRNA level using the following qPCR primers: GAPDH-fwd: ACTCCACTCACGGCAAATTCAACG, GAPDH-rev: AAGACACCAGTAGACTCCACGACA, beta-Actin-fwd: AGAGGGAAATCGTGCGTGAC, beta-Actin-rev: AACCGCTCGTTGCCAATAGT

### Microscopy

Cells were seeded on coverslips and transfected the next day using FuGENE6 (Promega) according to standard settings. The following day, cells were treated and immediately fixed with 4% PFA for 15 minutes at room temperature after washing twice with 1x PBS. Afterwards, cells on coverslips were washed three times with 1x PBS before dipping them in MilliQ water and drying them. After complete drying coverslips were mounted on a glass slide in 2.5µl DAPI-containing Vector Shield (Vector Laboratories) mounting medium and fixed on the slide to avoid dissolution. Glass slides were then kept at –20°C until use. Pictures were taken on a Leica DM4 B device using the LasX Software (Leica Microsystems).

### Bioinformatic analyses

The indicated data sets were analyzed using the Galaxy Suite (usegalaxy.org) (55). In brief, raw reads were downloaded from the GEO repository, demultiplexed and converted to the FASTQ format and groomed using “FASTQ Groomer“. After adapter- and quality-trimming using “Trim Galore!”, the reads were mapped against human hg19 or murine mm10 using “Bowtie2“. Aligned reads were then visualized in the Integrative Genomics Viewer (IGV) (56,57).

### Accession numbers

Nucleotide sequence data reported in this work is available in the Third Party Annotation (TPA) Section of the DDBJ/ENA/GenBank databases.

## Acknowledgement

The following people and institutions are thanked for providing expertise and reagents: Dr. Mathias Ballmaier and the sorting facility of the Hannover Medical School (MHH), Prof. Andreas Pich and the mass-spectrometry facility at the Hannover Medical School (MHH) for performing initial mass-spectrometry to identify isoforms. C.T. thanks Katrin Laaß for excellent technical assistance.

## Author Contributions

P.T. performed the majority of the experiments; M.B.M, G.S. and R.F. performed experiments; D.B.-J. and J.V.O. performed mass-spec experiments and the analyses; M.D.D.-M. performed bioinformatics analyses; M.G. and C.T. identified the odd reads in the 5’ UTR of the MK2 gene. A.K. designed experiments. M.T., M.G. and S.B.-J. supervised the study, provided lab space and contributed to the writing of the manuscript; C.T. performed experiments and wrote the paper.

All authors read and contributed to the editing of the final manuscript.

## Supplementary Figures

**Figure S1.** MK2 and MK3 expression and deletion in different murine and human cell lines. **A**. Detection of MK2, its phosporylated form (pMK2^T222^) and MK3 in MK2 KO and MK2/3 DKO bone marrow-derived macrophages (BMDMs) upon LPS-stimulation. The star indicates for a cross-reactive non-specific band detected by the MK2 antibody. Loading control: GAPDH **B**. Confirmation of the MK3-specific band in MK2 versus MK2/3 DKO MEFs by analyzing lysates of the indicted cells by Western Blot with a MK3-specific antibody. **C**. Detection of MK2 isoforms and MK3 in Jurkat-E6 and HCT116 cells by Western Blot using specific anti-MK2 and anit-MK3 antibodies. Loading control: GAPDH **D**. shRNA-mediated knockdown of MK2 in HT29 cells using different shRNAs raised against human MK2 (Particle #TRCN0000002282 and # TRCN0000002283) as well as a control shRNA (shCtrl) and magnitudes of infection (MOIs) were used. The efficiency of knockdown was analyzed by Western Blot of lysates probed with an MK2 antibody. Anti-GAPDH development served as a loading control.

**Figure S2.** Analysis of the human MK2 5′UTR sequence and its comparison to the mouse MK2 5′UTR. **A**. Genomic tracks for MK2 transcripts NM_004759.4 and NM_032960.3 in Ribosome Footprinting analyses derived from two different HEK293 data sets (GSE70804 and GSE37744). **B**. Human MK2 5′UTR nucleotide sequence and its putative extended in-frame N-terminal amino acid sequence. The aTIS are marked in bold and in italics within both nucleotide and the amino acid sequence. **C**. Alignment of the murine and human MK2 5′UTR nucleotide (lower) and amino acid (upper panel) sequences. The positions of putative aTIS are marked by red arrows within the amino acid sequence alignments.

**Figure S3.** Ribosome Footprinting analyses of murine bone marrow-derived dendritic cells (BMDCs) and keratinocytes. Upper panel (GSE74139): BMDCs were treated with cycloheximide (CHX), Lactimidomycin (LTM) or Harringtonine (Har) to freeze ribosomes on the mRNAs at different positions. Lower panel (GSE83332): Keatinocytes were either treated with cycloheximide (CHX) or Harringtonine (Har) to freeze ribosome on mRNAs in order to detect their positions form sequencing results. In both analyses the position of ribosomal footprints is shown for the 5′UTR of the murine MK2 transcript NM_008551.2.

**Figure S4.** Ribosome Footprinting analyses of HEK293, Jurkat and HCT116 cells. Upper panel: HEK293 (GSE37744) cells were treated with cycloheximide (CHX) or Harringtonine (Har; GSE59095) to freeze ribosomes on the mRNAs at different positions. Middle panel (GSE74279): Jurkat were either treated with cycloheximide (CHX) or Lactimidomycin (LTM) to freeze ribosome on mRNAs in order to detect their positions form sequencing results. Lower panel (GSE87328): HCT116 cells were treated with cycloheximide (CHX) or Lactimidomycin (LTM) to freeze ribosomes on mRNAs on specific positions. In all analyses the position of ribosomal footprints is shown for the 5′UTR of the human MK2 transcripts NM_004759.4 and NM_032960.3

**Figure S5.** MK2 5′UTR cloning strategy, introduction of mutation and their consequences after transfection in cells. **A**. Schematic representation of the MK2 5′UTR cloning strategy. The hitherto known murine MK2 coding sequence was cloned into peGFP-C1 allowing its expression in mammalian cells with a N-terminal GFP-fusion. By cutting with NheI (right after the CMV promotor of peGFP-C1) and PstI (within the known MK2 coding sequence), this part was replaced with a synthesized 5′UTR sequence containing the same sites at the 3’ and 5’ ends. This resulted in the elimination of the GFP-tag, but CMV-promotor driven expression of the complete 5′UTR in fusion with the MK2 coding sequence. This construct then contained the natural AUG translation initiation site and the other two potential aTIS CUG and GUG in position –141 and –309, respectively. The table below shows the different mutants that were derived from the initial construct #1 that contained the wildtype 5′UTR sequence. The predicted consequences of the mutations are given in the last column of the table. **B**. Expression of MK2 mutants in U2OS cells after transient transfection. The same plasmids as in Figure 2A were transfected into U2OS cells to prove effect the predicted consequences from the given in S5A. Lysates of transfected cells were analyzed by Western Blot with a MK2-specific antibody. Loading control: anti-p150.

**Figure S6.** MK2 translation mechanisms. **A**. Quantifications of band intensities from Fig. 2B using ImageJ. MK2 protein level were calculated in relation to GAPDH protein level. The line at 0.5 determines the approximate half-life of the MK2 isoforms in both genetic backgrounds. **B**. Schematic models of the MK2 truncation constructs used to analyze 5′UTR-dependent MK2 translation in Fig. 2C. **C**. Endogenous MK2 isoform translation upon eIF2A knockdown in RAW264.7, wildtype MEFs, HeLa and HEK293T cells. Cells were siRNA transfected with two different specific siRNAs or a control siRNA and lysates were subsequently analyzed by Western Blot for eIF2A to analyze knockdown efficiency and for MK2 to detect translation of isoforms. p150 served as loading control. **D**. Endogenous MK2 isoform translation upon Mettl3 knockdown in RAW264.7, wildtype MEFs, HeLa and HEK293T cells. Cells were siRNA transfected with two different Mettl3-specific siRNAs or a control siRNA and lysates were subsequently analyzed by Western Blot for Mettl3 to analyze knockdown efficiency and for MK2 to detect translation of isoforms. Tubulin served as loading control.

**Figure S7.** Expression of C-terminal GFP-tagged MK2 isoforms in HeLa cells. **A**. Transfected HeLa cells were analyzed for the expression GFP-tagged MK2 isoforms. The initial N-terminal tagged GFP-MK2 construct (peGFP-C1-MK2, see Fig. S5A) was used as a control as well as the peGFP-N1 vector used to clone the C-terminally tagged construct used in fluorescence microscopy. Western Blots were developed with anti-MK2- and anti-GFP-specific antibodies. Loading control: GAPDH. The translocation of MK2 isoforms from the nucleus to the cytosol after exposition to Anisomycin for 60 minutes was analyzed by fluorescence microscopy. U2OS cells were transfected with plasmids coding for C-terminal tagged isoforms of murine MK2. For this the mutant (construct #3 and #4, see Figure 2A and Figure S5) were subcloned into peGFP-N1 vectors that allow C-terminal GFP-tagging of inserted coding sequences. **B**. MEFs were analyzed for the phosphorylation of known MK2 substrates (NOGO-B and Hsp27) upon Anisomycin stimulation. Phosphorylation of the MK2 upstream p38^MAPK^ (pp38^T180/Y182^) and MK2 (pMK2^T334^) itself was used as a readout for pathway activation. To control activity, cells were either pre-incubated with an p38^MAPK^ inhibitor (Birb796) or a MK2 inhibitor (PF-3644022). p150 development served as loading control.

**Figure S8.** Functions of MK2 isoforms in vivo. **A**. Determining the growth of complemented MK2/3 DKO MEFs by a WST-1 assays. 1 × 10^4^ cells of every genotype were seeded in triplicates for each timepoint in a 96-well plate format. At the indicated times WST-1 was added to the cells and after 30 minutes of incubation at 37°C, densiometric measurements were performed in a multi-well plate reader. Relative OD600nm are plotted against the time [days]. **B**. Rescued MK2 MEFs were starved for 24h in serum-free DMEM and afterwards stimulated with 10% FCS-containing DMEM for 1 and 2h. Lysates were consequently analyzed for the expression of the indicated proteins. **C**. Expression of TTP mRNA in rescued MK2 MEFs after starvation as in Fig. S9B and stimulation with 10% FCS-containing DMEM for 0, 2 and 4h. Error bars represent the mean of three independent experiments (n = 3) +/− the standard deviation. P-values were retrieved from one-way ANOVA testing using the PRISM 7 software and are related to GFP/EV-rescued MEFs.

**Table S1.**
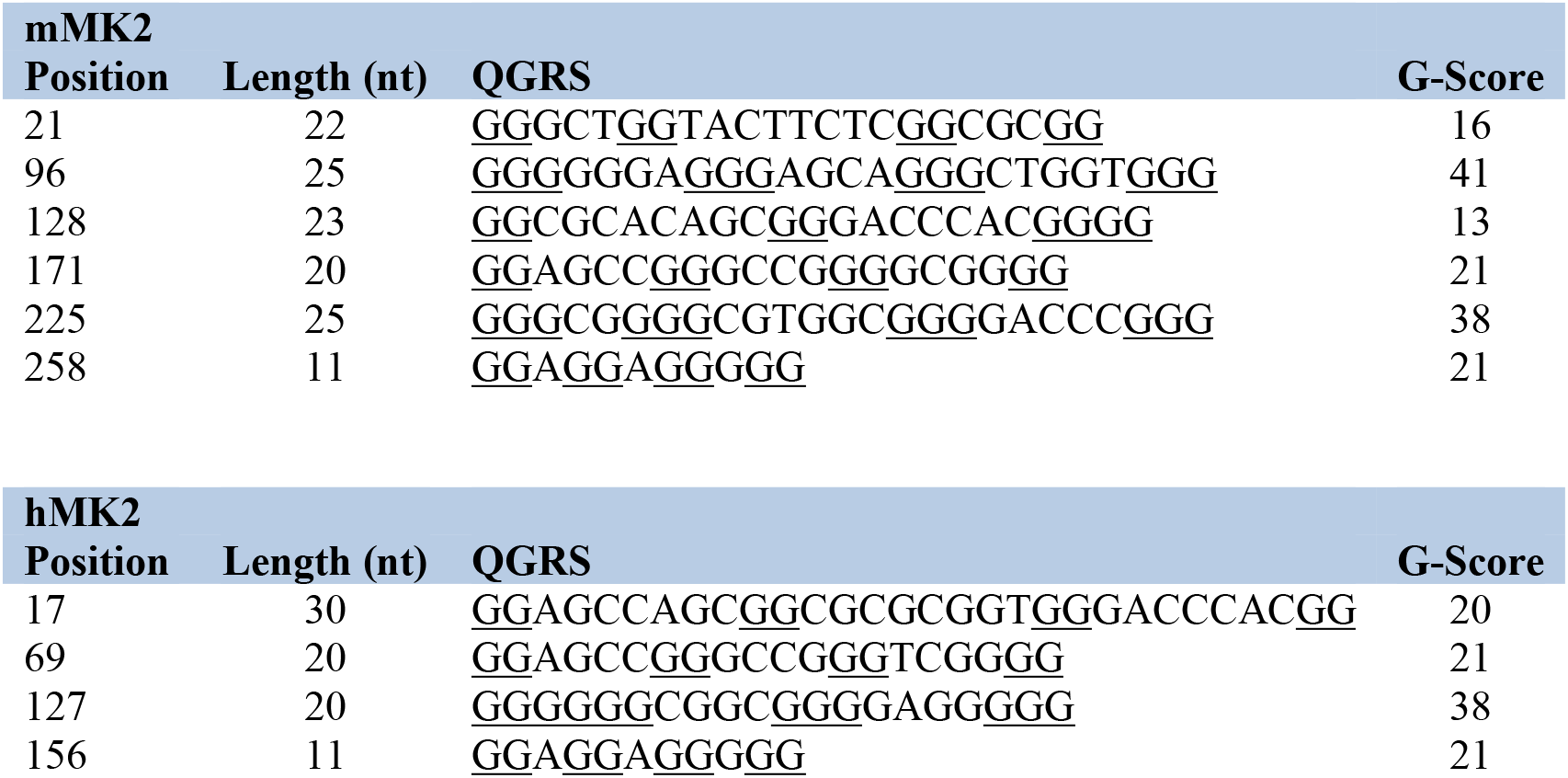
G-rich quadruplexes in GC-rich sequences in the MK2 5′UTR. Analysis of putative G-rich quadruplex forming sequences within the 5′UTR of murine and human MK2 using the QGRS online tool (1). High G-scores indicate for potential quadruplex structures. Under default settings, the highest possible G-score is 100.

